# A library of structurally homogeneous human *N*-glycans synthesized from microbial oligosaccharide precursors

**DOI:** 10.1101/118216

**Authors:** Brian S. Hamilton, Joshua D. Wilson, Marina A. Shumakovich, Adam C. Fisher, James C. Brooks, Alyssa Pontes, Radnaa Naran, Christian Heiss, Chao Gao, Robert Kardish, Jamie Heimburg-Molinaro, Parastoo Azadi, Richard D. Cummings, Judith H. Merritt, Matthew P. DeLisa

## Abstract

Synthesis of homogenous glycans in quantitative yields represents a major bottleneck to the production of molecular tools for glycoscience, such as glycan microarrays, affinity resins, and reference standards. Here, we describe a combined biological/enzymatic method termed bioenzymatic synthesis that is capable of efficiently converting microbially-derived precursor oligosaccharides into structurally uniform human-type N-glycans. Unlike starting material obtained by chemical synthesis or direct isolation from natural sources, which can be time consuming and costly to generate, bioenzymatic synthesis involves precursors derived from renewable sources including wild-type *Saccharomyces cerevisiae* glycoproteins and lipid-linked oligosaccharides from glycoengineered *Escherichia coli.* Following deglycosylation of these biosynthetic precursors, the resulting microbial oligosaccharides are subjected to a greatly simplified purification scheme followed by structural remodeling using commercially available and recombinantly produced glycosyltransferases including key *N-* acetylglucosaminyltransferases (*e.g.*, GnTI, GnTII, and GnTIV) involved in early remodeling of glycans in the mammalian glycosylation pathway. Using this approach, preparative quantities of hybrid and complex-type *N*-glycans including asymmetric multi-antennary structures were generated all without the need of a specialized skillset. Collectively, our results reveal bioenzymatic synthesis to be a user-friendly methodology for rapidly supplying homogeneous oligosaccharide structures that can be used to understand the human glycome and probe the biological roles of glycans in health and disease.

## Introduction

Carbohydrate chains, known as glycans, represent a vast source of biological diversity across all domains of life. They play a key role in almost every aspect of normal physiology, as well as in the etiology of nearly every major disease (Varki, 1993). Approximately 1% of the human genome is dedicated to the biosynthesis and diversification of glycans, and the majority of human proteins are thought to be post-translationally modified by glycans, a process known as glycosylation (Apweiler et al., 1999; Varki and Marth, 1995). Glycosylation can occur at several amino acid residues, most commonly asparagines (*N*-linked) and serines or threonines (*O*-linked). As conjugates to proteins, glycans add an additional information layer and have direct biological effects, ranging from stabilizing protein folds (Hebert et al., 2014; Helenius and Aebi, 2001; Imperiali and O’Connor, 1999) to signaling stem cell fate (Du and Yarema, 2010; Lanctot et al., 2007; Sampathkumar et al., 2006). Glycans also feature prominently in disease. For example, in the context of cancer, tumor cells commonly express glycans at atypical levels or with altered structural attributes (Adamczyk et al., 2012; Hakomori, 1985; Kim and Varki, 1997) and there is growing interest in developing pharmaceutical agents that target these molecules (Fuster and Esko, 2005). However, while the functional importance of glycosylation is well established, methods for mass-producing glycans and glycoconjugates are lagging (Sheridan, 2007; Walt et al., 2012). Such technological gaps are to be expected in light of the immense structural diversity and information density of glycans in nature (Werz et al., 2007), which far exceeds that of DNA or proteins. Unfortunately, this complexity, coupled with a scarcity of materials, has hindered production of glycomolecules which in turn has limited the availability of chemically defined oligosaccharides for use as standards in glycan structure determination (Marino et al., 2010; North et al., 2009), as probes to characterize glycan-binding proteins (Oyelaran and Gildersleeve, 2009; Rillahan and Paulson, 2011), and as ligands in affinity resins (Cummings and Etzler, 2009).

To furnish oligosaccharides for biological and structural studies, a variety of methods have been described that enable laboratory preparation of these important compounds. One such method involves the enzymatic or chemical deglycosylation of glycoproteins from natural sources (*e.g.*, hen egg yolk) followed by a series of purification steps (Kajihara et al., 2004; Verostek et al., 2000). While natural sources are a rich supplier of glycans, the chromatography, desalting, and concentration steps used to purify released glycans are time consuming, expensive, and typically yield only small quantities of closely related structures that are difficult to fractionate. For these reasons, chemical or chemo-enzymatic production of synthetic carbohydrates has emerged as a preferred route for preparing libraries of pure oligosaccharides (Boltje et al., 2009; Lepenies et al., 2010; Li et al., 2015; Maki et al., 2016; Palcic, 2011; Schmaltz et al., 2011; Wang et al., 2013). In the case of chemical synthesis, target oligosaccharides have been generated by either solution-phase or solid-phase methods. Solution-phase synthesis utilizes a one-pot strategy, wherein differences in the reactivity of glycosyl donors allow for ligation in the proper oligosaccharide sequence. In solid-phase synthesis, the reducing-end sugar is connected to a solid support and each additional sugar residue is added sequentially, after which the glycan is cleaved from the resin. While recent advances to these methods have greatly improved the number and diversity of glycans that can be produced chemically, including the addition of non-natural residues, this approach remains limited by the cost and scale of production (Blow, 2009; Rich and Withers, 2009) and remains very time consuming, especially when highly complex structures are targeted (Boltje et al., 2009). Many of these issues can be circumvented by chemo-enzymatic methods, whereby a synthetic oligosaccharide precursor is modified by a range of glycosyltransferases (GTs) to supply more complex derivatives (Palcic, 2011; Schmaltz et al., 2011). Chemo-enzymatic synthesis is not only more economical, but it enables efficient generation of extremely complex glycan structures, including asymmetrically branched *N*-glycans that have proven inaccessible to other existing synthesis technologies (Wang et al., 2013).

These methods notwithstanding, new glycan synthesis technologies for glycobiology and glycomedicine that are accessible, low-cost, user-friendly, robust, and adaptable are in high demand. Along these lines, here we describe a combined biological/enzymatic route that we call bioenzymatic synthesis that enables efficient, low-cost production of structurally homogeneous human-type *N*-glycans. One of the main difficulties to chemo-enzymatic or enzymatic synthesis of glycan libraries is the limited accessibility to oligosaccharides as starting material for diversification by GTs. The starting material is typically obtained by chemical synthesis or direct isolation from natural sources, which can be time consuming and costly as discussed above. To bridge this technology gap, bioenzymatic synthesis leverages precursor oligosaccharides that are derived in large quantities from two complementary microbial sources, either invertase glycoprotein from wild-type *Saccharomyces cerevisiae* or lipid-linked oligosaccharides (LLOs) from glycoengineered *Escherichia coli* (Valderrama-Rincon et al., 2012). Following deglycosylation of these biosynthetic precursors, the microbially-derived oligosaccharides are subjected to a greatly simplified purification scheme followed by structural remodeling using GTs, including several from the mammalian glycosylation pathway that are not commercially available. The power of this methodology was demonstrated by supplying preparative quantities of 20 different eukaryotic *N*-glycans including asymmetric multi-antennary structures, several of which have not previously been synthesized by chemical or chemoenzymatic routes.

## Results

### Preparation of human-type oligomannose glycan precursors

To generate a renewable supply of precursor oligosaccharides (see **Supplementary Table S1** for complete list with chemical structure), we investigated two complementary sources of *N*-glycans: hypermannoyslated invertase produced by recombinant expression in yeast; and undecaprenol pyrophosphate (Und-PP)-linked paucimannose glycans (*e.g.,* mannose_3_-*N*-acetylglucosamine_2_, a.k.a. Man_3_GlcNAc_2_) produced by glycoengineered *E. coli* (Valderrama-Rincon et al., 2012). Our initial focus was on invertase (Fig. 1a), which was chosen as a vehicle to isolate oligosaccharides because it has the highest glycan abundance of all known glycoproteins, and is produced at high yields when expressed recombinantly in wild-type yeast strains. Invertase carries high-mannose glycans comprised of eight or more mannose units (Trimble et al., 1991; Verostek et al., 2000); however, at the core of these glycans is a common intermediate structure, Man_5_GlcNAc_2_, that is also present in mammalian *N*-glycans. Previous work has engineered yeast to eliminate hypermannosylation, producing an oligosaccharide that is sensitive to mammalian glycosylation enzymes (Chiba et al., 1998; Hamilton et al., 2006; Okbazghi et al., 2016); however, to our knowledge, there has been no investigation of whether wild-type, hypermannosylated glycans can be converted to a glycoform that is sensitive to mammalian glycosylation enzymes. Thus, we investigated whether remodeling invertase glycans was a viable method for production of large amounts of starting material for production of a mammalian *N*-linked glycan library.

**Figure 1.**
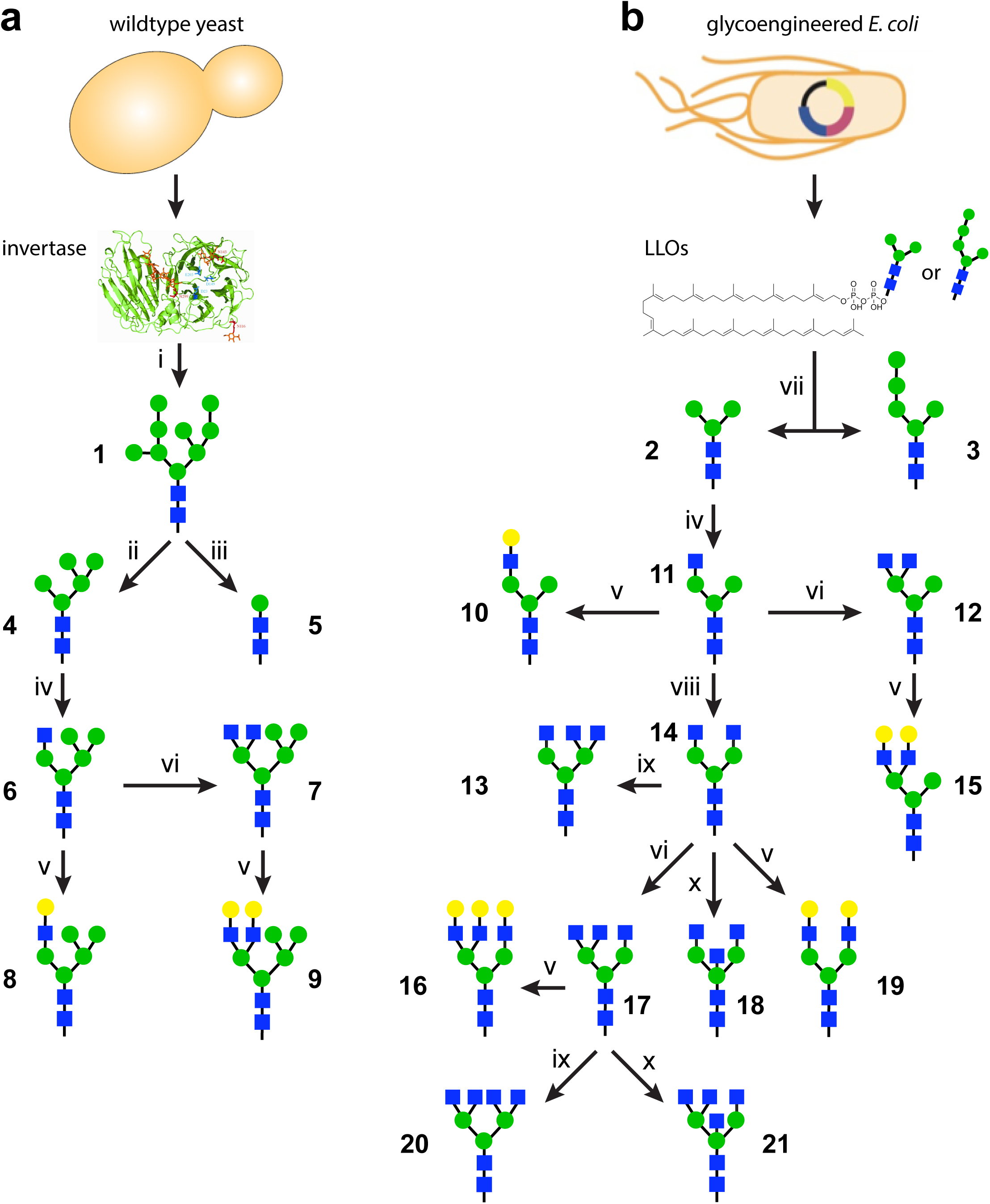
Glycan synthesis strategies. Precursor glycans (**1-3**) were derived from (a) yeast invertase or (b) lipid-linked oligosaccharides (LLOs) from glycoengineered *E. coli* cells carrying plasmid pYCG, and subsequently used to synthesize glycans **4-21**. Enzymatic steps: (i) PNGase F (product shown is representative of high mannose yeast glycans); (ii) α1,2- and α1,6- mannosidase; (iii) jack bean α-mannosidase and α1,6-mannosidase; (iv) GnTI; (v) β1,4- galactosyltransferase; (vi) GnTIV; (vii) non-enzymatic hydrolysis of extracted LLOs; (viii) GnTII; (ix) GnTV; and (x) GnTIII.

The glycan profile of the oligomannose glycans released from *Saccharomyces cerevisiae* invertase by PNGase F treatment revealed a mixture of Man_8_GlcNAc_2_ to Man_14_GlcNAc_2_ oligomannose glycans (**Supplementary Fig. S1a**). To determine whether it was possible to trim these high-mannose glycans to the human-type Man_5_GlcNAc_2_, we subjected released glycan 1 to a series of mannosidases and determined that the additional mannose residues consisted of α(1,2)- and α(1,6)-mannose linkages. By addition of *Xanthomonas manihotis* α(1,2)-mannosidase and *Aspergillus saitoi* α(1,6)-mannosidase, hypermannosylated yeast glycans were trimmed to the human-type Man_5_GlcNAc_2_ glycoform 4 whereas treatment with *Canavalia ensiformis* (jack bean) α-mannosidase and the same α(1,6)-mannosidase resulted in glycoform **5**. As discussed above, one of the drawbacks to isolation of glycans from natural sources is the difficulty in purification. Therefore, a point of emphasis was to develop a simplified purification method. After testing multiple separation techniques, we determined that an initial silica column followed by a charcoal/celite with a single elution step, and a graphitized carbon column with a gradient elution was the simplest method. This purification method resulted in a product having >95% purity as assessed by mass spectrometry (MS) and nuclear magnetic resonance (NMR) spectroscopy (Fig. 2a). Isolation, mannose trimming, and purification of the oligosaccharides from 1 g of invertase resulted in a yield of 0.7 mg of 4 as determined by measuring dry weight and fluorophore-assisted carbohydrate electrophoresis (FACE) analysis (Gao, 2005) (**Fig. S1b**). A similar procedure yielded glycan 5 (**Fig. S2a**). Thus, we confirm that biosynthesis of *S*. *cerevisiae* invertase glycans provides a viable reservoir of human-type oligosaccharide precursors.

**Figure 2.**
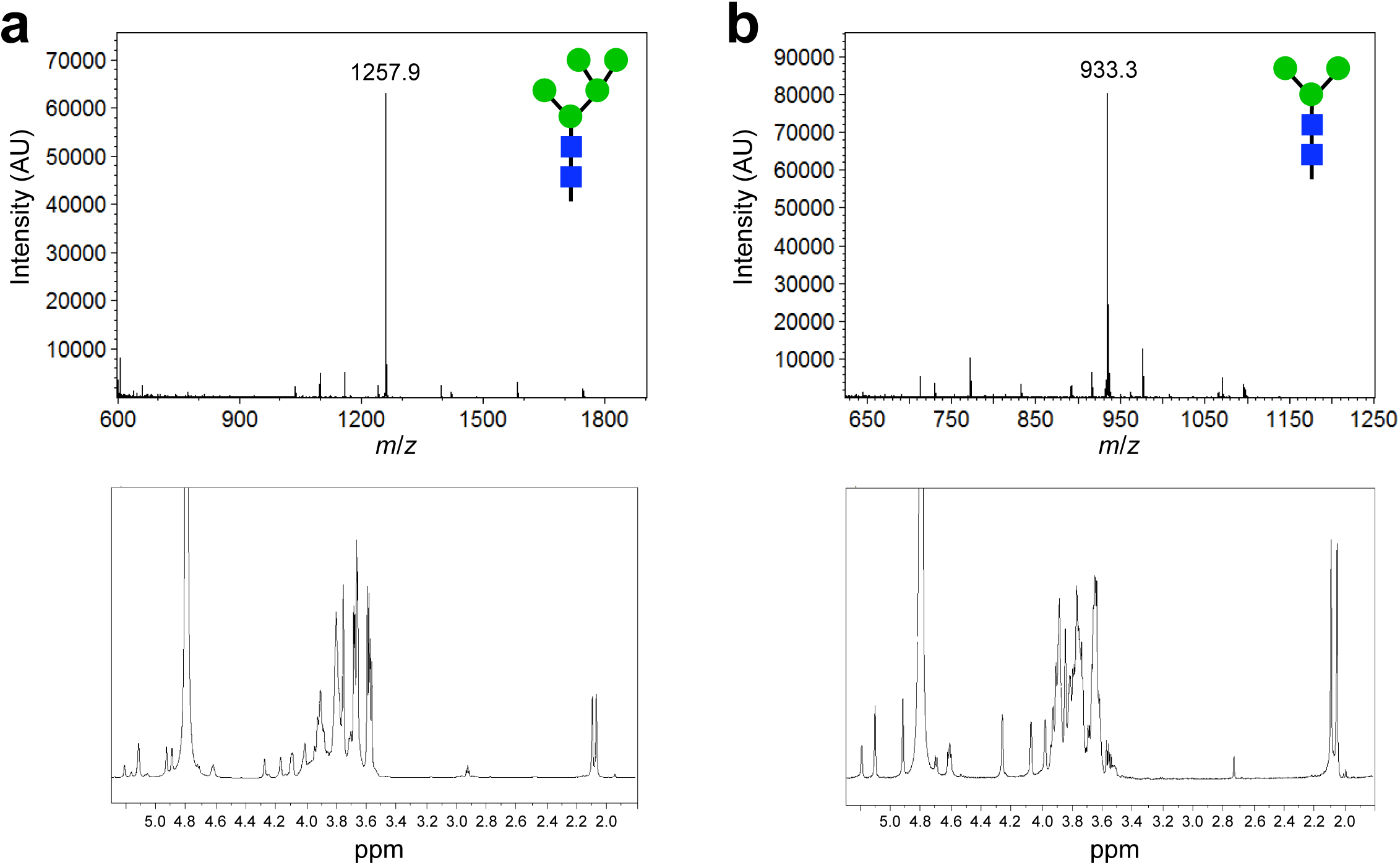
Biosynthesis of precursor oligosaccharides for enzymatic remodeling. MALDI-TOF MS analysis (top panels) and 600-MHz ^1^H NMR characterization (bottom panels) corresponding to: (a) Man_5_GlcNAc_2_ glycan synthesized by enzymatic deglycosylation of *S*. *cerevisiae* oligosaccharides; and (b) the Man_3_GlcNAc_2_ glycan synthesized by glycoengineered *E. coli* cells.

As an alternative biosynthetic route to precursor glycans, we investigated the use of glycoengineered *E. coli* as producers of the starting material for glycan diversification (Fig. 1b). Unlike all natural or engineered eukaryotic hosts, *E. coli* cells do not have native glycosylation pathways nor is glycosylation an essential mechanism in this host. As a result, *E. coli* provides an unformatted “operating system” that can be genetically reprogrammed to produce discrete, uniform glycans that are not attached to undesired targets and do not cause problems with cell physiology or morphology. Using natural pools of nucleotide sugars (*e.g.,* UDP-GlcNAc, GDP-Man) as substrates, *E. coli* cells carrying a synthetic pathway of heterologous GTs, namely the β1,4-GlcNAc transferase Alg13/14, the β1,4-mannosyltransferase Alg1, and the bifunctional mannosyltransferase Alg2, are capable of synthesizing Und-PP-linked Man_3_GlcNAc_2_ as we showed previously (Valderrama-Rincon et al., 2012). This latter product was readily isolated by lipid extraction from *E. coli* followed by non-enzymatic removal of **2** from undecaprenol, resulting in a structurally uniform Man_3_GlcNAc_2_ glycoform (Fig. 2b). It should be noted that addition of yeast mannosyltransferase Alg11 to the synthetic glycosylation pathway resulted in formation of Und-PP-linked **3** that could be similarly processed to generate nearly pure Man_5_GlcNAc_2_ glycans (**Fig. S2b**). Isolation of lipid-linked glycans from *E. coli* in this manner eliminates several difficulties associated with purification and heterogeneity. Moreover, the short time requirements and low costs associated with bacterial culture are advantageous for the synthesis of human-type *N*-glycan precursors.

### Expression and purification of eukaryotic glycosyltransferases

A number of the glycosylation enzymes that are natively involved in the early stages of remodeling oligomannose glycan **4** are not commercially available, and methods to produce these enzymes have not been reported. Therefore, we undertook the development of methods for recombinant expression and purification of three *N*-acetylglucosaminyltransferases (GnTs), namely GnTI, GnTII, and GnTIV. We chose *E. coli* due to its proven track record as a protein expression host. To promote efficient folding in the cytoplasm, the genes encoding *Nicotiana tabacum* GnTI, *Homo sapiens* GnTII, and *Bos taurus* GnTIV were all fused to the C-terminus of the gene encoding *E. coli* maltose binding protein (MBP) lacking its native export signal. Both GnTI and GnTIV were solubly expressed over a wide range of temperatures (15-37°C) and in a number of different *E. coli* strain backgrounds (*e.g.,* MC4100, SHuffle T7 Express, and Origami2(DE3)). In contrast, GnTII was only expressed solubly at temperatures at or below 25°C and required a host strain lacking both the thioredoxin reductase *(trxB)* and glutathione reductase (gor) genes (*e.g.,* Origami 2(DE3)), which greatly enhances disulfide bond formation in the *E. coli* cytoplasm (Bessette et al., 1999). All three enzymes were purified by amylose affinity chromatography and retained as fusions to MBP for enzymatic remodeling of yeast-derived glycan 4 and bacteria-derived glycan **2** as described below.

### Bioenzymatic synthesis of hybrid glycans from yeast-derived precursor

Hybrid oligosaccharide structures are inherently difficult to isolate from natural sources because they are transient glycans in the mammalian glycosylation pathway. Therefore, to produce these rare oligosaccharides, we conducted a 200-μg pilot scale synthesis of four target glycans using the yeast-derived Man_5_GlcNAc_2_ (glycan **4**) as the starting material. First, MBP-GnTI was used to generate a hybrid GlcNAcMan_5_GlcNAc_2_ glycan **6** by conjugating a β1,2-linked GlcNAc residue to the α1,3 arm of **4**. Production of this hybrid glycan was confirmed by MS and NMR analysis (Fig. 3a). Importantly, the chemical shifts of **6** synthesized herein matched previous NMR characterization of this oligosaccharide structure (Table 1) (Chen et al., 2008). One advantage of enzymatic glycan synthesis is the ability to conduct multiple reactions in a single step. Synthesis of **7** is a good example of this, where both MBP-GnTI and MBP-GnTIV were added simultaneously to produce a glycan with β1,4-GlcNAc branching on the α1,3 mannose (Fig. 3b). Conducting multiple reactions in a single step eliminates purification of each intermediate, which in turn reduces loss of product associated with purification. As an example of further elaboration of these oligosaccharides, each glycan was treated with *B. taurus* β1,4-galactosyltransferase to produce glycans **8** and **9**. Addition of galactose residues was confirmed by MS, and by the appearance of a peak that corresponds to galactose in each NMR spectrum (Fig. 3c and d). In the case of glycan **9**, the reaction must be conducted in a two-step process to eliminate the possibility of producing **8** as a side product. Taken together, these results confirm bioenzymatic synthesis of human-type, hybrid oligosaccharides.

**Table 1.** Chemical shift assignments of the synthesized N-linked oligosaccharides.

|  | 2 | 3 | 4 | 5 | 6 | 7 | 8 | 9 | 10 | 11 | 12 | 13 | 14 | 15 | 16 | 17 | 18 | 19 | 20 | 21 |
| --- | --- | --- | --- | --- | --- | --- | --- | --- | --- | --- | --- | --- | --- | --- | --- | --- | --- | --- | --- | --- |
| H-1 |  |  |  |  |  |  |  |  |  |  |  |  |  |  |  |  |  |  |  |  |
| GlcNAc-1α | 5.21 | 5.20 | 5.21 | 5.21 | 5.19 | 5.20 | 5.20 | 5.21 | 5.21 | 5.20 | 5.20 | 5.21 | 5.20 | 5.20 | 5.20 | 5.20 | 5.20 | 5.21 | 5.21 | 5.21 |
| GlcNAc-1β | 4.72 | 4.71 | 4.72 | 4.72 | 4.70 | 4.71 | 4.71 | 4.71 | 4.72 | 4.70 | 4.72 | 4.71 | 4.70 | 4.70 | 4.70 | 4.70 | 4.71 | 4.71 | 4.71 | 4.71 |
| GlcNAc-2 | 4.63 | 4.61 | 4.62 | 4.62 | 4.60 | 4.61 | 4.60 | 4.61 | 4.63 | 4.62 | 4.62 | 4.61 | 4.61 | 4.61 | 4.61 | 4.61 | 4.62 | 4.63 | 4.61 | 4.62 |
| Man-3 | 4.80 | 4.80 | 4.80 | 4.81 | 4.80 | 4.80 | 4.80 | 4.80 | 4.80 | 4.80 | 4.80 | 4.80 | 4.80 | 4.80 | 4.80 | 4.80 | 4.71 | 4.80 | 4.80 | 4.71 |
| Man-4 | 5.12 | 5.35 | 5.11 | ----- | 5.13 | 5.12 | 5.13 | 5.13 | 5.12 | 5.13 | 5.13 | 5.12 | 5.12 | 5.12 | 5.12 | 5.12 | 5.08 | 5.13 | 5.13 | 5.07 |
| Man-4' | 4.92 | 4.92 | 4.89 | ----- | 4.89 | 4.89 | 4.89 | 4.89 | 4.92 | 4.93 | 4.93 | 4.88 | 4.93 | 4.92 | 4.93 | 4.93 | 5.02 | 4.93 | 4.88 | 5.02 |
| Man-A | ----- | ----- | 5.11 | ----- | 5.11 | 5.10 | 5.11 | 5.11 | ----- | ----- | ----- | ----- | ----- | ----- | ----- | ----- | ----- | ----- | ----- | ----- |
| Man-B | ----- | ----- | 4.93 | ----- | 4.93 | 4.92 | 4.93 | 4.93 | ----- | ----- | ----- | ----- | ----- | ----- | ----- | ----- | ----- | ----- | ----- | ----- |
| Man-C | ----- | 5.31 | ----- | ----- | ----- | ----- | ----- | ----- | ----- | ----- | ----- | ----- | ----- | ----- | ----- | ----- | ----- | ----- | ----- | ----- |
| Man-D | ----- | 5.05 | ----- | ----- | ----- | ----- | ----- | ----- | ----- | ----- | ----- | ----- | ----- | ----- | ----- | ----- | ----- | ----- | ----- | ----- |
| GlcNAc-5 | ----- | ----- | ----- | ----- | 4.58 | 4.57 | 4.59 | 4.59 | 4.59 | 4.57 | 4.57 | 4.57 | 4.56 | 4.56 | 4.58 | 4.54 | 4.57 | 4.58 | 4.57 | 4.54 |
| GlcNAc-5' | ----- | ----- | ----- | ----- | ----- | 4.55 | ----- | 4.57 | ----- | ----- | 4.54 | 4.57 | 4.56 | 4.56 | 4.58 | 4.54 | 4.57 | 4.58 | 4.57 | 4.57 |
| GlcNAc-6 | ----- | ----- | ----- | ----- | ----- | ----- | ----- | ----- | ----- | ----- | ----- | ----- | ----- | ----- | 4.58 | 4.54 | ----- | ----- | 4.54 | 4.57 |
| GlcNAc-6' | ----- | ----- | ----- | ----- | ----- | ----- | ----- | ----- | ----- | ----- | ----- | 4.57 | ----- | ----- | ----- | ----- | ----- | ----- | 4.54 | ----- |
| GlcNAc-7 | ----- | ----- | ----- | ----- | ----- | ----- | ----- | ----- | ----- | ----- | ----- | ----- | ----- | ----- | ----- | ----- | 4.48 | ----- | ----- | 4.48 |
| Gal-8 | ----- | ----- | ----- | ----- | ----- | ----- | 4.49 | 4.49 | 4.48 | ----- | ----- | ----- | ----- | 4.47 | 4.48 | ----- | ----- | 4.49 | ----- | ----- |
| Gal-8' | ----- | ----- | ----- | ----- | ----- | ----- | ----- | 4.48 | ----- | ----- | ----- | ----- | ----- | 4.47 | 4.48 | ----- | ----- | 4.49 | ----- | ----- |
| Gal-9 | ----- | ----- | ----- | ----- | ----- | ----- | ----- | ----- | ----- | ----- | ----- | ----- | ----- | ----- | 4.48 | ----- | ----- | ----- | ----- | ----- |
| NHAc |  |  |  |  |  |  |  |  |  |  |  |  |  |  |  |  |  |  |  |  |
| CH<sub>3</sub>CO | 2.05 | 2.06 | 2.06 | 2.06 | 2.06 | 2.07 | 2.07 | 2.07 | 2.06 | 2.06 | 2.06 | 2.06 | 2.06 | 2.05 | 2.06 | 2.06 | 2.06 | 2.06 | 2.06 | 2.06 |
| CH<sub>3</sub>CO | 2.09 | 2.09 | 2.09 | 2.09 | 2.09 | 2.08 | 2.08 | 2.08 | 2.08 | 2.08 | 2.08 | 2.09 | 2.08 | 2.08 | 2.08 | 2.09 | 2.09 | 2.08 | 2.09 | 2.08 |
| CH<sub>3</sub>CO | ----- | ----- | ----- | ----- | 2.06 | 2.05 | 2.05 | 2.06 | 2.05 | 2.05 | 2.05 | 2.05 | 2.05 | 2.04 | 2.05 | 2.05 | 2.05 | 2.05 | 2.05 | 2.05 |
| CH<sub>3</sub>CO | ----- | ----- | ----- | ----- | ----- | 2.10 | ----- | 2.10 | ----- | ----- | 2.10 | 2.09 | 2.05 | 2.10 | 2.05 | 2.09 | 2.04 | 2.05 | 2.09 | 2.04 |
| CH<sub>3</sub>CO | ----- | ----- | ----- | ----- | ----- | ----- | ----- | ----- | ----- | ----- | ----- | 2.06 | ----- | ----- | 2.09 | 2.06 | 2.09 | ----- | 2.06 | 2.09 |
| CH<sub>3</sub>CO | ----- | ----- | ----- | ----- | ----- | ----- | ----- | ----- | ----- | ----- | ----- | ----- | ----- | ----- | ----- | ----- | 2.07 | ----- | 2.04 | ----- |

**Figure 3.**
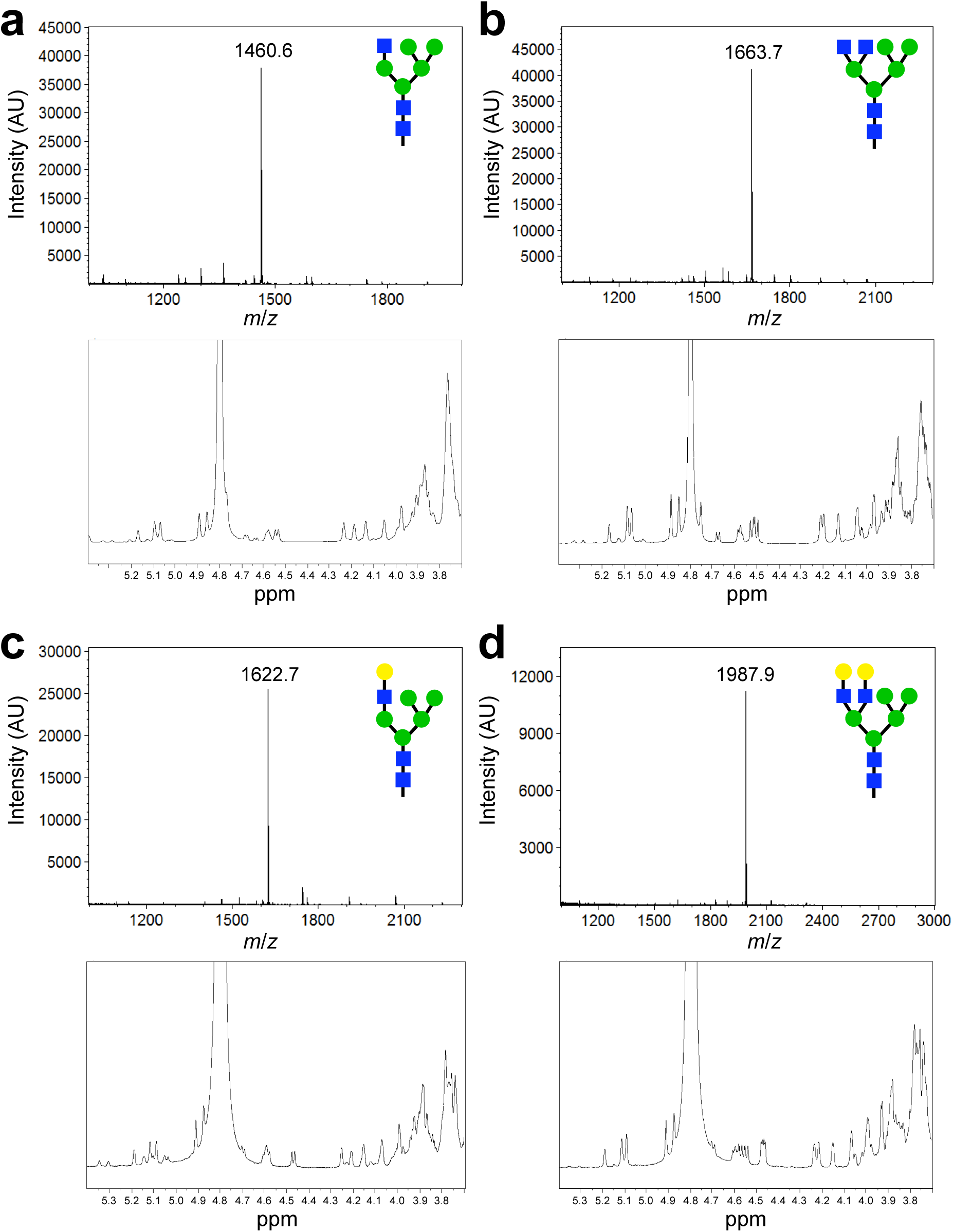
Bioenzymatic synthesis of hybrid glycans using the yeast-derived precursor. MALDI-TOF MS analysis (top panels) and 600-MHz ^1^H NMR characterization (bottom panels) for the following: (a) glycan **6**; (b) glycan **7**; (c) glycan **8**; and (d) glycan **9**.

### Bioenzymatic synthesis of hybrid and complex glycans from bacteria-derived precursor

We next evaluated whether the *E. coli-derived* Man_3_GlcNAc_2_ glycan **2** could be used as the starting material for 200-μg pilot-scale synthesis of hybrid and complex-type glycans. An advantage of this precursor glycan is that it eliminates two mannosidase-catalyzed deglycosylation steps required to produce the hybrid GlcNAcMan_3_GlcNAc_2_ glycan **11**. Indeed, following treatment with MBP-GnTI, efficient transfer of a GlcNAc residue directly to the α1,3 arm of **2** was observed (Fig. 4a), despite the fact that this precursor glycan is not the native substrate for this enzyme. The chemical shifts of glycan **11** were in agreement with previous NMR spectroscopy of this oligosaccharide structure (Table 1) (Kajihara et al., 2004). Along similar lines, we synthesized the human-type complex GlcNAc_2_Man_3_GlcNAc_2_ glycan **14** using a one-pot strategy whereby both MBP-GnTI and MBP-GnTII were added simultaneously to **2** (Fig. 4b). This glycan was also characterized by NMR and matched previous characterization of this oligosaccharide structure (Table 1) (Kajihara et al., 2004). It is worth mentioning that the resulting complex glycan is a key intermediate in the mammalian *N*-glycosylation pathway, and in our glycan library synthesis efforts, because it acts as a substrate for all subsequent additions. For example, further GlcNAc addition was possible by treatment of **11** or **14** with MBP-GnTIV to produce glycans **12** and **17**, respectively, with β1,4-GlcNAc branching (Fig. 4c and d, and Table 1). Interestingly, when carrying out the synthesis of glycan **17**, the reaction rate of GnTII was greatly decreased when acting on glycan **12** in the one-pot reaction. Therefore, glycan **17** was synthesized in a stepwise fashion whereby a first reaction with MBP-GnTI and MBP-GnTII was followed by a second reaction with MBP-GnTIV. Glycans with β1,6-GlcNAc branching were achieved by step-wise treatment of **14** or **17** with *H. sapiens* GnTV to yield **13** and **20**, respectively (Fig. 4e and f, and Table 1). Another interesting point is that two pairs of the GlcNAc-terminal glycans **12** and **14**, **13** and **17** are structural isomers, which would be useful as standards for glycomics research and as affinity probes for determining lectin substrate specificity. Collectively, these results demonstrate that treatment of the bacteria-derived precursor glycan with different combinations of GnTI, GnTII, GnTIV, and GnTV is an efficient route to multiantennary, GlcNAc-terminal oligosaccharides.

**Figure 4.**
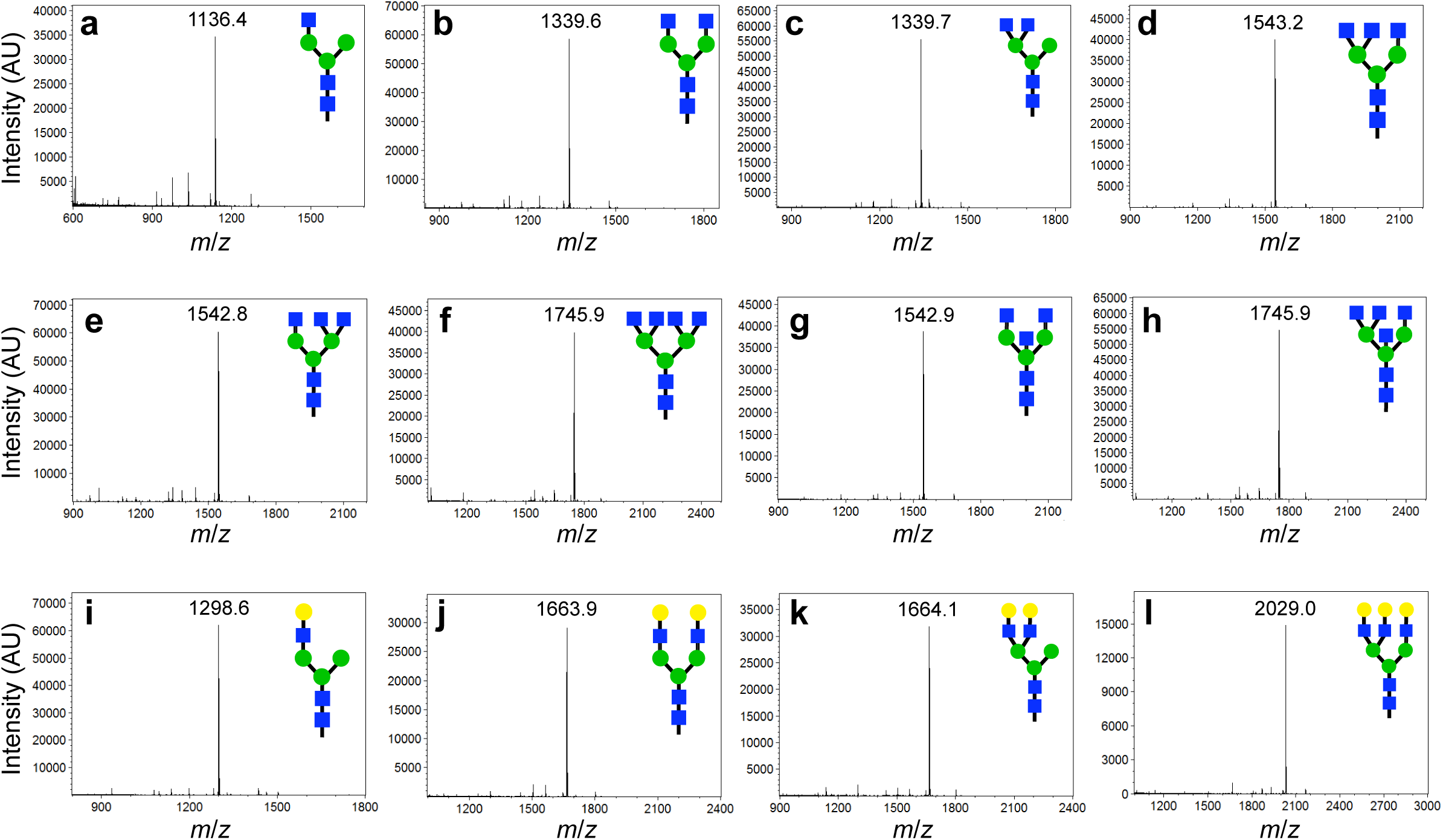
Bioenzymatic synthesis of hybrid and complex-type glycans using the bacteria-derived precursor. MALDI-TOF MS analysis of the following products: (a) glycan **11**; (b) glycan **14**; (c) glycan **12**; (d) glycan **17**; (e) glycan **13**; (f) glycan **20**; (g) glycan **18**; (h) glycan **21**; (i) glycan **10**; (j) glycan **19**; (k) glycan **15**; (l) glycan **16**.

Additional modification of select oligosaccharides described above was conducted to demonstrate the rapid increase in the number of glycan structures that can be produced by our bioenzymatic method. For example, *H. sapiens* GnTIII adds a bisecting GlcNAc to the β1,4-mannose and is thought to be a natural regulator of branching, since attachment of this residue eliminates further elaboration by GnTs. To generate such bisecting glycan structures, glycans **14** and **17** were synthesized in a first reaction and then, for reasons described above, treated in a subsequent reaction with GnTIII to yield **18** and **21**, respectively. Analysis of the reaction products revealed that GnTIII installed branching GlcNAc residues at a high conversion rate (Fig. 4g and h, and Table 1). The resulting bisecting GlcNAc structures, which supply additional structural isomers to our panel, would be of particular interest for determining differences in lectin binding specificity. Finally, galactose residues were successfully added to **11**, **12**, **14**, and **17** by treatment with *B. taurus* β1,4-galastosyltransferase, yielding glycans **10**, **15**, **19**, and **16**, respectively (Fig. 4i-l, and Table 1). While these latter reactions were only conducted on select glycan structures with two different GTs, one could easily apply these GTs to each glycan produced herein, resulting in an exponential increase in the number of oligosaccharides in the glycan library.

### Construction and evaluation of an *N*-glycan microarray

To demonstrate the functionality of the glycan library, we developed microarrays using a novel bifunctional fluorescent linker, 2-amino-*N*-(2-aminoethyl)-benzamide (AEAB) (Song et al., 2009). Specifically, glycans **2-21** were directly conjugated to AEAB through its arylamine group by reductive amination to form glycan-AEABs (GAEABs) and then purified by multidimensional HPLC. Following purification, GAEABs were covalently immobilized onto *N*-hydroxysuccinimide (NHS)-activated glass slides via their free alkylamine alongside the similarly immobilized reference compounds LNnT, LNT, and NA2 (Fig. 5a), which were prepared from natural glycans as described previously (Song et al., 2009). When probed with *C. ensiformis* Concanavalin A (ConA), binding of the biantennary *N*-glycan reference compound, NA2, was detected but not reference compounds LNT or LNnT, consistent with the specificity of ConA for primarily internal and non-reducing terminal α-mannose (Fig. 5b). Among the bioenzymatic glycans, ConA bound most strongly to the oligomannose and hybrid glycans (**2-12**). It also bound complex glycans **13**, **14**, **17**, **18**, **20**, and **21**, albeit to a lesser extent, which was consistent with the established higher affinity of ConA for terminal versus internal α-linked mannose. In agreement with previous array data (http://www.functionalglycomics.org/glycomics/HServlet?operation=view&sideMenu=no&psId=primscreen_PA_v2_357_10312005), ConA bound efficiently to the complex glycan 19, which is structurally identical to reference compound NA2. In contrast, ConA showed virtually no binding to its hybrid isomer, glycan **15**. Taken together, these results demonstrate the compatibility of bioenzymatically-derived glycans with microarray development (*i.e.,* chemical derivatization and immobilization steps) and fluorescence-based screening of glycan-binding proteins.

**Figure 5.**
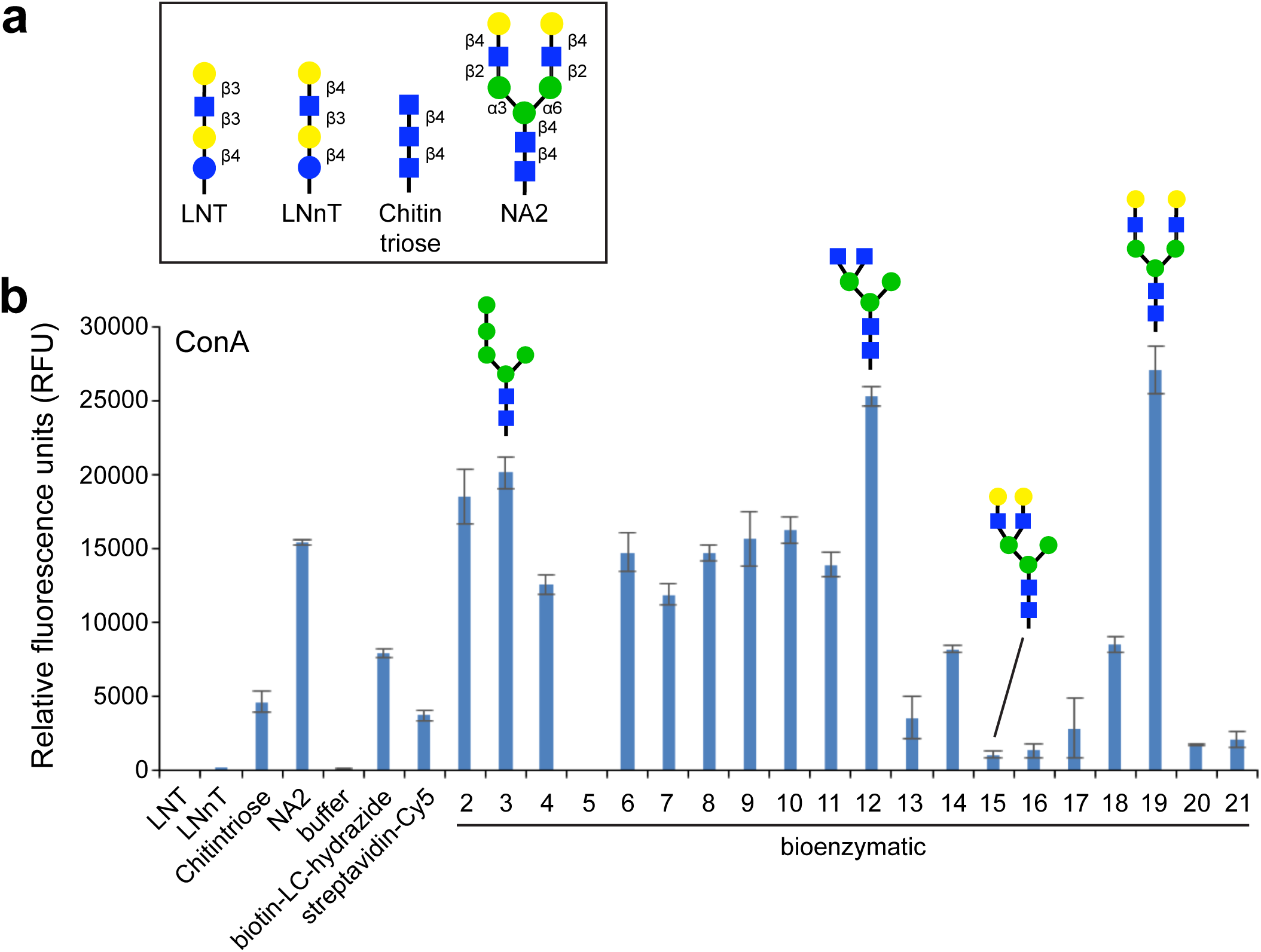
Binding of lectin ConA to microarray of bioenzymatically produced *N*-glycans. (a) Structures of reference compounds that were derivatized with AEAB and immobilized onto NHS-activated glass slides, alongside glycans **2-21**. (b) Probing of immobilized glycans with biotinylated ConA (10 μg/ml). The amount of bound ConA was determined by streptavidin-Cy5 (5 μg/ml) fluorescence. Background subtracted mean fluorescence values are shown. Error bars represent the standard deviation of the mean. Representative structures for glycans **3**, **12**, **15** and **19** are shown. All glycans were printed at 100 μM in PBS in replicates of four. PBS and streptavidin-Cy5 spots served as controls. Biotin-LC-hydrazide spots printed on each subarray serve as an alignment feature to localize each subarray on the slide.

## Discussion

As many have argued (Walt et al., 2012), the development of simple and cost-effective methods to produce diverse glycomolecules is needed to meet the demand of, and grow, the glycoscience community. In this work, we addressed this need with the demonstration of an efficient new methodology – called bioenzymatic synthesis – that enables production of large quantities of hybrid-and complex-type *N*-glycans. This was accomplished by first harnessing the power of microbial biosynthesis to furnish a renewable reservoir of starting material. Specifically, up to milligram amounts of precursor oligosaccharides were derived from either wild-type yeast that recombinantly express hypermannosylated invertase or glycoengineered *E. coli* that produce lipid-linked paucimannose glycans. In parallel, we developed simplified methods to purify the resulting oligosaccharides, as well as robust expression and purification methods for key glycan remodeling enzymes (*e.g.,* eukaryotic GnTs) that are not commercially available. Armed with these and other commercially available GTs, we efficiently transformed precursor oligosaccharides into a library of hybrid and complex *N*-glycans including asymmetric multi-antennary structures, all in appreciable yields and without the need of a specialized skillset. We anticipate that this technology will help to overcome many of the critical barriers to glycan production by offering a user-friendly synthesis methodology that can reliably deliver homogeneous glycans from a consistent source and flexibly adapt to production needs. Moreover, the libraries of glycans that can be supplied by the bioenzymatic methodology presented here represent useful tools for understanding the biological role of glycans in health and disease as well as for defining and exploiting the human glycome.

## Experimental Procedures

### Strains and plasmids

*E. coli* strain MC4100 (F^−^ *araD139* ∆(*argF-lac*)*U169 flbB5301 deoC1 ptsF25 relA1 rbsR22 rpsL150 thiA*), SHuffle T7 Express (New England Biolabs), and Origami2(DE3) (Novagen) were used for heterologous expression of different eukaryotic GnTs.

Origami2(DE3) gmd::kan *∆waaL ∆nanA* cells were used for producing lipid-linked Man_3_GlcNAc_2_ precursor. This strain was created by introducing sequential mutations using P1 *vir* phage transduction (Thomason et al., 2007) with the respective strains from the Keio collection (Baba et al., 2006) as donors. Briefly, to delete the *nanA* gene a lysate was generated from donor cells (JW31194-1) containing the *∆nanA735::kan* mutation. Donor strains were obtained from the Coli Genetic Stock Center (CGSC). The resulting phage was used to infect Origami2(DE3) target cells and successful transductants were selected on LB plates supplemented with kanamycin (Kan). The Kan resistance cassette was subsequently removed by transforming the resulting strain with plasmid pCP20 as described (Datsenko and Wanner, 2000). This resulted in generation of strain Origami2(DE3) *∆nanA.* Subsequent deletion of the *waaL* (*rfaL*) and *gmd* genes was performed in the same manner using donor strains JW3597-1 (Δ*rfaL734::kan)* and JW2038-1 (*∆*gmd751::kan). Yeast strain FY834 was used for homologous recombination using plasmid pMQ70 as described previously (Shanks et al., 2006). YPD growth medium was used to maintain yeast and synthetic defined uracil dropout medium was used to select and maintain yeast with plasmids. Plasmid pYCG was used for production of lipid-linked Man_3_GlcNAc_2_ and has been described previously (Valderrama-Rincon et al., 2012). A second plasmid containing the *E. coli manB* and *manC* genes for over-production of GDP-mannose was produced using homologous recombination in *S*. *cerevisiae* (Shanks et al., 2006). Briefly, genes encoding phosphomannomutase (ManB) and mannose-1-phosphate guanylyltransferase (ManC) were PCR-amplified from *E. coli* genomic DNA using primer pairs that contained the priming region for the *manB* and *manC* genes, respectively, along with 40-bp overhangs of pMQ70. The resulting PCR products and the linearized pMQ70 plasmid were used to transform yeast strain FY834, yielding plasmid pManCB. Constructs assembled in yeast were electroporated into *E. coli* MC4100 for verification via PCR, restriction enzyme digestion, and/or sequencing. For expression of eukaryotic GnTs, *N. tabacum* GnTI, *H. sapiens* GnTII, *B. taurus* GnTIV were all cloned for expression as C-terminal fusions to *E. coli* MBP lacking its native export signal peptide. Briefly, genes encoding MBP and the respective GnT enzyme were amplified with primers containing regions of overlap to the fusion partner and pMQ70 plasmid. The PCR products were assembled with linearized pMQ70 by homologous recombination as described above, yielding plasmids pMBP-GnTI, pMBP-GnTII, and pMBP-GnTIV.

### Synthesis of Man_5_GlcNAc_2_ precursor

5 g of *S*. *cerevisiae* invertase (Sigma-Aldrich) was denatured in the presence of sodium dodecyl sulfate and NP-40 as described in the New England Biolabs protocol for PNGase F treatment. The solution was allowed to cool to room temperature and 50,000 U of PNGase F (New England Biolabs) was added to the solution. The PNGase F reaction was incubated at 30°C for 2 days to ensure complete removal of oligosaccharide from the protein. The glycan was precipitated using 80% (v/v) acetone, and resuspended in 60% methanol (v/v), where the remaining pellet was discarded. The glycan/protein mixture was then applied to a water-equilibrated charcoal/celite (1:1) column and eluted with 50% ethanol (v/v). The glycan was then dried by rotary evaporator, resuspended in a minimal amount of ethyl acetate:methanol:water (11:3:3), and applied to a 11:3:3 pre-equilibrated silica flash column. The column was washed with a ratio of 7:3:3 and eluted with a ratio of 3:3:3. The glycan was then resuspended in water and desalted by mixing with Ag 1-X8 and DOWEX 50WX8-400 resins. Both the characterization and purity were determined by MALDI-TOF MS, FACE, and 600-MHz 1D ^1^H NMR spectroscopy as described elsewhere (Gao, 2005; Kajihara et al., 2004; Valderrama-Rincon et al., 2012). The resulting high-mannose precursor oligosaccharides were specifically trimmed with both X. *manihotis* α1-2-mannosidase (ProZyme) and *A. saitoi* α1-6-mannosidase (New England Biolabs) to produce the human-type Man_5_GlcNAc_2_ glycan, since we empirically determined that there were no terminal α(1,3)-mannose residues. Alternatively, to generate the Man_1_GlcNAc_2_ glycan, *C. ensiformis* α-mannosidase (Sigma) and *A. saitoi* α(1,6)-mannosidase were used to trim invertase-derived glycans. To achieve this conversion, roughly 5 mg of purified precursor glycan was resuspended in mannosidase buffer (0.1 M sodium acetate, pH 5.0, ProZyme), to which 8 mU of a1-2-mannosidase and 80 U of α1-6-mannosidase were added. The reaction was allowed to proceed at 37°C and monitored by MS until the reaction was complete. The glycan was then purified by a graphitized carbon column (Hypercarb, Thermo) using a 0-50% acetonitrile:water gradient. Characterization and purity of the glycan was assessed by 600-MHz NMR.

### Synthesis of Man_3_GlcNAc_2_ precursor

Origami2(DE3) *gmd::kan ∆nanA ∆waaL* cells were transformed with plasmids pYCG and pManCB by electroporation and used to inoculate a shake flask containing 200 mL LB containing 2% (v/v) glucose along with 100 μg/mL ampicillin (Amp) and 25 μg/mL chloramphenicol (Cam), and incubated overnight, with shaking at 30°C. 50 mL of the 200-mL culture was then transferred to each of eight 1-L shake flasks containing LB supplemented with Amp and Cam and cultures were incubated overnight, with shaking at 30°C. The cells were then pelleted by centrifugation at 4,236 × g, resuspended in 5 mL of methanol, and sonicated five times at 10-sec pulses. The sonicated lysate was then poured into a glass petri dish and heated at 60°C until dry. The methods for extraction of LLOs and release of the glycan from the lipid were followed as described in Gao et al. (Gao, 2005). Briefly, *E. coli* cell pellets were resuspended in 2:1 chloroform:methanol, sonicated, and the remaining solids collected by centrifugation. This pellet was sonicated in water and collected by centrifugation. The resulting pellet was sonicated in 10:10:3 chloroform:methanol:water to isolate the LLOs from the inner membrane. The LLOs were purified using acetate-converted DEAE anion exchange chromatography as they bind to the anion exchange resin via the phosphates that link the lipid and glycan. The resulting compound was dried and treated by mild acid hydrolysis to release glycans from the lipids. The released glycans were then separated from the lipid by a 1:1 butanol:water extraction, wherein the water layer contains the glycans. The glycans were then further purified with a graphitized carbon column using a 0-50% water:acetonitrile gradient. The Man_3_GlcNAc_2_ glycan was analyzed by MADLI-TOF MS, FACE, and 600-MHz ^1^H NMR.

### Expression and purification of eukaryotic GnTs

*E. coli* strains MC4100, SHuffle, or Origami2(DE3) were separately transformed with pMBP-GnTI, pMBP-GnTII, or pMBP-GnTIV by electroporation. Transformed cells were transferred to starter cultures containing 50 mL of LB with 2% (v/v) glucose and 100 μg/mL Amp and incubated overnight at 30°C. The 50-mL cultures were then transferred to a shake flask containing 1 L LB supplemented with 100 μg/mL Amp and induced for 16 h with 0.2% arabinose when the absorbance (Abs_600_) reached 1.0. Cells were harvested by centrifugation at 4,320 x g, resuspended in a minimal amount of MBP binding buffer (20 mM Tris-HCl, 1 mM EDTA, 200 mM NaCl, pH 7.4), and sonicated 5 times at 30-sec pulses. The lysate was subjected to centrifugation at 12,000 x g for 20 min, and the cell debris was discarded. The clarified lysate was loaded onto an MBP binding buffer, preequilibrated amylose gravity flow column (New England Biolabs) and washed with the binding buffer. The protein was eluted with 10 mL MBP binding buffer supplemented with 10 mM maltose and concentrated to 1 mL by centrifugation (Vivaspin 20, GE Healthcare).

### Synthesis of hybrid oligosaccharides from the yeast-derived precursor

To produce 6, a 500-μg sample of **4** was treated with an excess amount of GnTI and UDP-GlcNAc (Sigma-Aldrich) in GnT buffer (20 mM HEPES, 50 mM NaCl, 10 mM MnSO_4_, pH 7.2), and incubated at 37°C until the reaction was complete, as determined by MS. To synthesize **7**, a 500-μg sample of **4** was treated with an excess amount of GnTI and GnTIV along with an excess of UDP-GlcNAc and incubated at 37°C until the reaction was complete. Each glycan was desalted as described above and purified by silica flash column chromatography as described above. The glycans were then characterized by MALDI-TOF MS and 1D ^1^H 600-MHz NMR. A 250-μg sample of **6** and **7** were treated with an excess amount of *B. taurus* β1,4-galactosyltransferase (Sigma-Aldrich) and UDP-Gal (Sigma-Aldrich) to produce **8** and **9**. The reaction was carried out at 37°C until completion, as determined by MS. The glycan products were purified and analyzed described above.

### Synthesis of hybrid and complex glycans using the bacteria-derived precursor

Ten 1-L cultures of Man_3_GlcNAc_2_-producing *E. coli* cells were required for pilot-scale synthesis of each glycan. A ~250-μg sample of Man_3_GlcNAc_2_ was used for synthesis of each glycan produced. Both UDP-GlcNAc and UDP-Gal were added in 3-fold molar excess, and each enzyme was also added in excess as determined empirically. All synthesized glycans were desalted as described above and purified by graphitized carbon using a 0-50% acetonitrile:water gradient. The purified products were analyzed by MALDI-TOF MS, FACE, and 600-MHz ^1^H NMR.

### Synthesis of GlcNAc terminal glycans

To produce glycan **11**, Man_3_GlcNAc_2_ was treated with GnTI and incubated overnight at 37°C. To produce **14**, both GnTI and GnTII were added simultaneously to **2** in GnT buffer and incubated overnight at 37°C. To synthesize **12**, GnTI and GnTIV were added simultaneously to **2** in GnT buffer and incubated overnight at 37°C. For synthesis of **13**, GnTI, GnTIV, and *H. sapiens* GnTV (R&D systems) were added simultaneously to **2** in GnTI buffer and incubated at 37°C until the reaction was complete. To synthesize **17**, GnTI and GnTII were added simultaneously to **2** in GnT buffer and incubated at 37°C until the reaction was complete. GnTIV was then added to the solution and incubated at 37°C until the reaction was complete. Similarly, **20** was synthesized by adding GnTI and GnTII to **2** in GnT buffer and incubated at 37°C until the reaction was complete. GnTIV and GnTV were then added to the solution and incubated at 37°C until the reaction was complete. The progress of each reaction was monitored by MALDI-TOF MS and determined to be complete when the substrate glycan was undetectable.

### Synthesis of bisecting GlcNAc glycans

To produce **18** and **21**, the glycans **14** and **17** were first synthesized as described above, and used as substrates for the addition of bisecting GlcNAc. *H. sapiens* GnTIII (R&D systems) was added to **14** and **17** separately in GnT buffer and incubated at 37°C until the reaction was complete, as determined by MS.

### Synthesis of Gal terminal glycans

For synthesis of **10**, GnTI and GalT were added simultaneously to **2** in GnTI buffer and incubated at 37°C until the reaction was complete. For synthesis of **15**, **16**, and **19**, the GlcNAc terminal glycans (**12**, **14**, **17**) were first synthesized as described above before the addition of Gal to avoid production of **10** as a side product. GalT was added to **12**, **14**, and **17** in GnT buffer and incubated at 37°C until the reactions were complete as determined by MS.

### NMR analysis

Glycans 2, 3, 5, 10-21 (≥100 μg each, Complex Carbohydrate Research Center, University of Georgia, acquired spectra and performed analysis) were deuterium exchanged 3 times by suspending in D_2_O and subsequent lyophilization, were re-dissolved in 300 μl D_2_O (99.96%, Cambridge Isotopes) and placed in a 3-mm Shigemi tubes. 1-D proton (2-D gCOSY, zTOCSY, and 2D-HSQC were also acquired as needed) spectra were obtained at 25°C on Varian Inova 600 MHz spectrometer equipped with cryoprobe using standard Varian pulse sequences. Glycans 4, and 6-9 (≥ 100 μg each) were prepared as above, except rotary evaporation was used for deuterium exchange (D_2_O, 99,9%, Sigma), and 1-D proton spectra were acquired at 25°C using a Varian Inova 600-MHz spectrometer (Cornell University) with a pulse field gradient probe. Chemical shifts were measured relative to HOD peak (δ_H_=4.82 ppm).

### Glycan-AEAB conjugation and purification

Free reducing glycans were conjugated with AEAB as described previously (Song et al., 2009). Briefly, 25 μg of glycan was mixed with 10 μl of AEAB hydrochloride salt solution freshly prepared at 0.5 M in DMSO/AcOH (7:3, v/v) followed by an equal volume of 1.0 M sodium cyanoborohydride solution freshly prepared in the same solvent. The mixture was vortexed briefly and incubated at 65°C for 4 h. The mixture was chilled, and the products were precipitated upon the addition of 10 volumes of acetonitrile. After bringing the suspension to −20°C and maintaining that temperature for 2 h, the products were separated by centrifugation at 10,000 × *g* for 3 min. The pellet was dried by centri-vap and stored at −20°C for further purification. AEAB derivatives of LNnT, LNT, and the biantennary *N*-glycan, NA2, were prepared as reference standards.

### Solid phase extraction by-NH_2_ column

The dried sample was reconstituted in 10 μl of water, loaded onto the −NH_2_ SPE column, which had been prewashed with acetonitrile, water and 85% acetonitrile in water. The −NH_2_ SPE column was then washed with 3 bed volumes of acetonitrile, followed by 3 bed volumes of 85% acetonitrile and the AEAB conjugate was eluted by 5% MeCN, 100 mM ammonium acetate. The combined eluents were dried under vacuum to remove the acetonitrile and lyophilized repeatedly to remove the ammonium acetate.

### Printing, binding assay, and scanning

NHS-activated slides were purchased from Schott (Louisville, KY). Non-contact printing was performed using a Scienion printer. The printing, binding assay, and scanning conditions were essentially the same as described previously (Song et al., 2011). Briefly, all samples were printed at 100 μM in phosphate buffer (300 mM sodium phosphates, pH 8.5) in replicates of four. Before the assay, the slides were rehydrated for 5 min in TSM wash buffer (20 mM Tris-HCl, 150 mM sodium chloride (NaCl), 2 mM calcium chloride (CaCl_2_), and 2 mM magnesium chloride (MgCl_2_)). Samples were applied to the rehydrated slide in TSM buffer containing 1% BSA and 0.05% Tween 20 in a final volume of 100 μl and the slides were incubated in a humidified chamber at room temperature for 1 h. After incubation, sample was washed away by gently dipping the slides in buffer contained in Coplin jars. Biotinylated ConA (Vector Labs) lectin binding to glycans on the array was detected by a secondary incubation with streptavidin-cyanine 5 at 5 μg/ml (SA-Cy5). The slides were scanned with a Genepix fluorescence microarray scanner equipped with four lasers covering an excitation range from 488 to 637 nm. For Cy5 fluorescence, 633 nm (excitation) was used at laser power 70% and PMT 450. All images obtained from the scanner are in *grayscale* and *colored* for easy discrimination. The scanned images were analyzed and quantitated in relative fluorescence units (RFU) from each spot in an Excel spreadsheet, to determine the average and S.D. of the four replicates. The %CV (coefficient of variation expressed in percent) was calculated as 100 × S.D./mean.

### Data availability

All datasets related to glycan array screening that were generated in this work are available through The Functional Glycomics Gateway (http://www.functionalglycomics.org/glycomics/publicdata/home.jsp), a comprehensive and free online resource provided by the CFG.

## Acknowledgements

This research was supported by National Science Foundation Awards CBET-1605242, CBET-1159581, and MCB-1411715 (to M.P.D.), the New York State Office of Science, Technology and Academic Research Distinguished Faculty Award (to M.P.D.), the National Institutes of Health Small Business Innovation Research grants 1R43GM088905-01 and 2R44GM088905-02 (to J.H. M.), and the National Institutes of Health contract HHSN261201300046C (to J.H.M.). The NMR analysis was performed at the Complex Carbohydrate Research Center, supported in part by a grant from the NIH-funded Research Resource for Integrated Glycotechnology (5P41GM10339024 to P.A.). The glycan microarray analysis was performed at the National Center for Functional Glycomics (NCFG) at Beth Israel Deaconess Medical Center, supported in part by the National Center for Research Resources (P41GM103694 to R.D.C.). We also thank Anthony Condo and Ivan Keresztes of the Chemistry NMR Facility (Cornell University) for their assistance acquiring NMR spectra.

## References

Adamczyk, B., Tharmalingam, T., and Rudd, P.M. (2012). Glycans as cancer biomarkers. Biochim Biophys Acta 1820, 1347–1353.

Apweiler, R., Hermjakob, H., and Sharon, N. (1999). On the frequency of protein glycosylation, as deduced from analysis of the SWISS-PROT database. Biochim Biophys Acta 1473, 4–8.

Baba, T., Ara, T., Hasegawa, M., Takai, Y., Okumura, Y., Baba, M., Datsenko, K.A., Tomita, M., Wanner, B.L., and Mori, H. (2006). Construction of Escherichia coli K-12 in-frame, singlegene knockout mutants: the Keio collection. Mol Syst Biol 2, 2006–0008.

Bessette, P.H., Aslund, F., Beckwith, J., and Georgiou, G. (1999). Efficient folding of proteins with multiple disulfide bonds in the Escherichia coli cytoplasm. Proc Natl Acad Sci U S A 96, 13703–13708.

Blow, N. (2009). Glycobiology: A spoonful of sugar. Nature 457, 617–620.

Boltje, T.J., Buskas, T., and Boons, G.J. (2009). Opportunities and challenges in synthetic oligosaccharide and glycoconjugate research. Nat Chem 1, 611–622.

Chen, R., Pawlicki, M.A., Hamilton, B.S., and Tolbert, T.J. (2008). Enzyme-Catalyzed Synthesis of a Hybrid N-Linked Oligosaccharide using N-Acetylglucosaminyltransferase I. Advanced Synthesis & Catalysis 350, 1689–1695.

Chiba, Y., Suzuki, M., Yoshida, S., Yoshida, A., Ikenaga, H., Takeuchi, M., Jigami, Y., and Ichishima, E. (1998). Production of human compatible high mannose-type (Man5GlcNAc2) sugar chains in Saccharomyces cerevisiae. J Biol Chem 273, 26298–26304.

Cummings, R.D., and Etzler, M.E. (2009). Antibodies and lectins in glycan analysis. In Essentials of Glycobiology (Cold Spring Harbor (NY): Cold Spring Harbor Laboratory Press).

Datsenko, K.A., and Wanner, B.L. (2000). One-step inactivation of chromosomal genes in Escherichia coli K-12 using PCR products. Proc Natl Acad Sci U S A 97, 6640–6645.

Du, J., and Yarema, K.J. (2010). Carbohydrate engineered cells for regenerative medicine. Adv Drug Deliv Rev 62, 671–682.

Fuster, M.M., and Esko, J.D. (2005). The sweet and sour of cancer: glycans as novel therapeutic targets. Nat Rev Cancer 5, 526–542.

Gao, N. (2005). Fluorophore-assisted carbohydrate electrophoresis: a sensitive and accurate method for the direct analysis of dolichol pyrophosphate-linked oligosaccharides in cell cultures and tissues. Methods 35, 323–327.

Hakomori, S. (1985). Aberrant glycosylation in cancer cell membranes as focused on glycolipids: overview and perspectives. Cancer Res 45, 2405–2414.

Hamilton, S.R., Davidson, R.C., Sethuraman, N., Nett, J.H., Jiang, Y., Rios, S., Bobrowicz, P., Stadheim, T.A., Li, H., Choi, B.K., et al. (2006). Humanization of yeast to produce complex terminally sialylated glycoproteins. Science 313, 1441–1443.

Hebert, D.N., Lamriben, L., Powers, E.T., and Kelly, J.W. (2014). The intrinsic and extrinsic effects of N-linked glycans on glycoproteostasis. Nat Chem Biol 10, 902–910.

Helenius, A., and Aebi, M. (2001). Intracellular functions of N-linked glycans. Science 291, 2364–2369.

Imperiali, B., and O’Connor, S.E. (1999). Effect of N-linked glycosylation on glycopeptide and glycoprotein structure. Curr Opin Chem Biol 3, 643–649.

Kajihara, Y., Suzuki, Y., Yamamoto, N., Sasaki, K., Sakakibara, T., and Juneja, L.R. (2004). Prompt chemoenzymatic synthesis of diverse complex-type oligosaccharides and its application to the solid-phase synthesis of a glycopeptide with Asn-linked sialyl-undeca- and asialo-nonasaccharides. Chemistry 10, 971–985.

Kim, Y.J., and Varki, A. (1997). Perspectives on the significance of altered glycosylation of glycoproteins in cancer. Glycoconj J 14, 569–576.

Lanctot, P.M., Gage, F.H., and Varki, A.P. (2007). The glycans of stem cells. Curr Opin Chem Biol 11, 373–380.

Lepenies, B., Yin, J., and Seeberger, P.H. (2010). Applications of synthetic carbohydrates to chemical biology. Curr Opin Chem Biol 14, 404–411.

Li, L., Liu, Y., Ma, C., Qu, J., Calderon, A.D., Wu, B., Wei, N., Wang, X., Guo, Y., Xiao, Z., et al. (2015). Efficient Chemoenzymatic Synthesis of an N-glycan Isomer Library. Chem Sci 6, 5652–5661.

Maki, Y., Okamoto, R., Izumi, M., Murase, T., and Kajihara, Y. (2016). Semisynthesis of Intact Complex-Type Triantennary Oligosaccharides from a Biantennary Oligosaccharide Isolated from a Natural Source by Selective Chemical and Enzymatic Glycosylation. J Am Chem Soc 138, 3461–3468.

Marino, K., Bones, J., Kattla, J.J., and Rudd, P.M. (2010). A systematic approach to protein glycosylation analysis: a path through the maze. Nat Chem Biol 6, 713–723.

North, S.J., Hitchen, P.G., Haslam, S.M., and Dell, A. (2009). Mass spectrometry in the analysis of N-linked and O-linked glycans. Curr Opin Struct Biol 19, 498–506.

Okbazghi, S.Z., More, A.S., White, D.R., Duan, S., Shah, I.S., Joshi, S.B., Middaugh, C.R., Volkin, D.B., and Tolbert, T.J. (2016). Production, Characterization, and Biological Evaluation of Well-Defined IgG1 Fc Glycoforms as a Model System for Biosimilarity Analysis. J Pharm Sci 105, 559–574.

Oyelaran, O., and Gildersleeve, J.C. (2009). Glycan arrays: recent advances and future challenges. Curr Opin Chem Biol 13, 406–413.

Palcic, M.M. (2011). Glycosyltransferases as biocatalysts. Curr Opin Chem Biol 15, 226–233.

Rich, J.R., and Withers, S.G. (2009). Emerging methods for the production of homogeneous human glycoproteins. Nat Chem Biol 5, 206–215.

Rillahan, C.D., and Paulson, J.C. (2011). Glycan microarrays for decoding the glycome. Annu Rev Biochem 80, 797–823.

Sampathkumar, S.G., Li, A.V., Jones, M.B., Sun, Z., and Yarema, K.J. (2006). Metabolic installation of thiols into sialic acid modulates adhesion and stem cell biology. Nat Chem Biol 2, 149–152.

Schmaltz, R.M., Hanson, S.R., and Wong, C.H. (2011). Enzymes in the synthesis of glycoconjugates. Chem Rev 111, 4259–4307.

Shanks, R.M., Caiazza, N.C., Hinsa, S.M., Toutain, C.M., and O’Toole, G.A. (2006). Saccharomyces cerevisiae-based molecular tool kit for manipulation of genes from gramnegative bacteria. Appl Environ Microbiol 72, 5027–5036.

Sheridan, C. (2007). Commercial interest grows in glycan analysis. Nat Biotechnol 25, 145–146.

Song, X., Lasanajak, Y., Xia, B., Heimburg-Molinaro, J., Rhea, J.M., Ju, H., Zhao, C., Molinaro, R.J., Cummings, R.D., and Smith, D.F. (2011). Shotgun glycomics: a microarray strategy for functional glycomics. Nat Methods 8, 85–90.

Song, X., Xia, B., Stowell, S.R., Lasanajak, Y., Smith, D.F., and Cummings, R.D. (2009). Novel fluorescent glycan microarray strategy reveals ligands for galectins. Chem Biol 16, 36–47.

Thomason, L.C., Costantino, N., and Court, D.L. (2007). E. coli genome manipulation by P1 transduction. Curr Protoc Mol Biol Chapter 1, Unit 1 17.

Trimble, R.B., Atkinson, P.H., Tschopp, J.F., Townsend, R.R., and Maley, F. (1991). Structure of oligosaccharides on Saccharomyces SUC2 invertase secreted by the methylotrophic yeast Pichia pastoris. J Biol Chem 266, 22807–22817.

Valderrama-Rincon, J.D., Fisher, A.C., Merritt, J.H., Fan, Y.Y., Reading, C.A., Chhiba, K., Heiss, C., Azadi, P., Aebi, M., and DeLisa, M.P. (2012). An engineered eukaryotic protein glycosylation pathway in Escherichia coli. Nat Chem Biol 8, 434–436.

Varki, A. (1993). Biological roles of oligosaccharides: all of the theories are correct. Glycobiology 3, 97–130.

Varki, A., and Marth, J. (1995). Oligosaccharides in vertebrate development. Seminars in Developmental Biology 6, 127–138.

Verostek, M.F., Lubowski, C., and Trimble, R.B. (2000). Selective organic precipitation/extraction of released N-glycans following large-scale enzymatic deglycosylation of glycoproteins. Anal Biochem 278, 111–122.

Walt, D., Aoki-Kinoshita, K.F., Bendiak, B., Bertozzi, C.R., Boons, G.-J., Darvill, A., Hart, G., Kiessling, L.L., Lowe, J., Moon, R.J., et al. (2012). Transforming Glycoscience: A Roadmap for the Future (The National Academies Press).

Wang, Z., Chinoy, Z.S., Ambre, S.G., Peng, W., McBride, R., de Vries, R.P., Glushka, J., Paulson, J.C., and Boons, G.J. (2013). A general strategy for the chemoenzymatic synthesis of asymmetrically branched N-glycans. Science 341, 379–383.

Werz, D.B., Ranzinger, R., Herget, S., Adibekian, A., von der Lieth, C.W., and Seeberger, P.H. (2007). Exploring the structural diversity of mammalian carbohydrates ("glycospace") by statistical databank analysis. ACS Chem Biol 2, 685–691.

